# Rapid Detection of VanA/B-Producing Vancomycin-Resistant enterococci using Lateral Flow Immunoassay from colonies and blood culture

**DOI:** 10.1101/2020.11.23.395293

**Authors:** Saoussen Oueslati, Hervé Volland, Vincent Cattoir, Sandrine Bernabeu, Delphine Girlich, Duncan Dulac, Marc Plaisance, Laure Boutigny, Laurent Dortet, Stéphanie Simon, Thierry Naas

## Abstract

Vancomycin-resistant enterococci (VRE) have become one of the most important nosocomial pathogens worldwide associated with increased treatment costs, prolonged hospital stay, and high mortality. Rapid detection is crucial to reduce their spread and prevent infections and outbreaks. The lateral flow immunoassay NG-Test VanB (NG Biotech) was evaluated for the rapid detection of VanB-producing vancomycin-resistant enterococci (VanB-VRE) using 104 well-characterized enterococcal isolates. The sensitivity and specificity were both 100%, when bacterial cells were grown in the presence of vancomycin used as VanB inducer. The NG-Test VanB is an efficient, rapid, and easy to implement assay in clinical microbiology laboratories for the confirmation of VanB-VREs either from colonies or from positive blood cultures. Together with the NG-Test VanA, they could complete/replace the already existing panel of tests available for confirmation of acquired vancomycin resistance in enterococci, especially from selective media (rectal screenings) or from antibiograms (infections), with a sensitivity and specificity of both of 100%. The rapid detection in less than 15 minutes will result in more efficient management of carriers and infected patients.

## Importance

Vancomycin-resistant enterococci (VRE) are increasingly isolated worldwide and constitute a major public health concern. Even though not very virulent, infections of hospitalized patients with VREs may be difficult to treat and, occasionally, life threatening. VRE is usually spread by direct contact with hands, environmental surfaces or medical equipment that has been contaminated by the feces of an infected person. Rapid detection of VRE is crucial for implementing appropriate infection control measures and to adapt antibiotic treatment for optimizing care strategies and outcomes. Lateral flow immunoassays (LFIAs) have shown their usefulness as easy, rapid and reliable confirmatory tests for antibiotic resistance mechanisms detection. We have validated an easy to implement LFIA, the NG-Test VanB, for the rapid (<15 min) identification of VanB-VRE from colonies or from positive blood cultures. The NG-Test VanB, combined with the NG-Test VanA allow identification of VanA and VanB-VREs, the two main acquired-vancomycin resistance determinants worldwide.

Vancomycin-resistant enterococci (VRE) are increasingly isolated worldwide and constitute a major public health concern (1-4). Vancomycin resistance is due to the expression of *van* operons. Currently, there are eight different acquired vancomycin resistance operons described: *vanA, vanB, vanD, vanE, vanG, vanL, vanM* and *vanN* (4-6). In Europe, the two main resistance phenotypes are by far VanA, and VanB. *Enterococci* expressing VanA (VanA-VREs) display high levels of inducible resistance to both vancomycin and teicoplanin, whereas strains expressing VanB (VanB-VRE) have variable levels of inducible resistance to vancomycin only (6,7). Rapid detection of VRE is crucial to help implement appropriate infection control measures and to adapt antibiotic treatment for optimizing care strategies and outcomes (4,8). Several culture-based methods, such as chromogenic or selective screening media, have been developed for VRE detection from rectal swabs. These methods usually take 24-48h, and as their specificity are low, colonies grown on these selective media require confirmatory testing, such as PCR (8,9,10). In addition, they may also lack sensitivity, especially with VanB-VREs with MICs of 4-8 µg/mL (4). Molecular techniques are much faster, as compared to culture, as they may be used directly on rectal swabs, but positive predictive value may also be low, especially for *vanB* genes that may be harbored by anaerobic bacteria part of the intestinal microbiota, and culture remains mandatory to confirm every *vanB*-positive result (10,11). Recently, the NG-Test VanA lateral flow immunoassay (LFIA) wa shown to be efficient for the detection of VanA-VRE from bacterial cultures (12). Here, we have developed a rapid and reliable companion LFIA, the NG-Test VanB for the detection of the second most prevalent VRE, *i*.*e*. VanB-VRE.

The *vanB* gene of *E. faecalis* V583 (4) was PCR amplified using the primers VanB NdeI (5’-gatataCATATGaatagaataaaagttgcaatactg3’) and VanB XhoI (5’-gtggtgCTCGAGcccctttaacgctaatacgatcaa3’) and then cloned into pET22b+ vector (Novagen; Merk, Darmstadt, Germany) (12-14). Recombinant plasmids pET22b+ *vanB* and pET22b-plyV12 (12,15), which encodes a broadly active phage lytic enzyme with lethal activity against *E. faecalis* and *E. faecium* were transformed into *Escherichia coli* BL 21 (DE3). Upon induction, the recombinant VanB and plyV12 proteins were purified using Ni-NTA agarose affinity resin as previously described (12-14). The VanB recombinant protein was then used to immunize mice and as a standard for the selection of monoclonal antibody (mAb) pairs as previously described (12-14). All animal experiments were performed in compliance with French and European regulations on the care of laboratory animals (European Community Directive 86/609, French Law 2001-486, 6 June 2001) and with the agreements of the Ethics Committee of the Commissariat à l’Energie Atomique (CEtEA) no. 12-026 and 15-055 delivered by the French Veterinary Services and CEA agreement D-91-272-106 from the Veterinary Inspection Department of Essonne (France). Ten-weeks-old Biozzi mice were immunized by intraperitoneal injection of purified recombinant VanB protein (50 µg), and the best pair of antibodies were produced on a large scale and provided to NG Biotech (Guipry, France) for the development of the NG-Test VanB assay, as previously described (12-14)

The detection of VanB-producers was first investigated using eight enterococcal isolates (3 VanB, 1 VanA, 1 VanC1, 1 VanC2 and 2 non-VRE) grown on different culture media widely used in routine: nine agar plates (Mueller-Hinton [MH], MH + a vancomycin disc of 5 µg, UriSelect™ 4, Bile Esculin Azide were from Bio-Rad [Marne-la-Coquette, France], ChromID^®^VRE, Columbia agar + 5% horse blood, Chocolate agar PolyViteX, and D-Coccosel agar were from bioMérieux [Marcy-l-Etoile, France] and an in-house prepared MH agar plate supplemented with 6 µg/ml of vancomycin) and two broth (brain heart infusion (BHI, bioMérieux) with and without a 30 µg vancomycin-containing disk. For bacterial colonies, 1-µL loop full of bacteria grown on the different agar plates was added to 100 µL extraction buffer (EB, provided by NG Biotech supplemented with 80 µg/mL of lysin (EB-80). For bacterial broth culture, 500 µL culture was centrifuged for 5 min at 10,000 rpm, the pellet was resuspended in 100 µL of EB-80 (12). After an incubation of 5 min at room temperature, the extract was loaded onto the cassette. The result was eye read after 15 min of migration by monitoring the appearance of a red band specific for VanB (test line, T), along with a band corresponding to the internal control (control line, C).

The NG-Test VanB was able to detect all VanB-VREs only when vancomycin was added into the medium, allowing induction of the *vanB* operon (Table 1, Figure 1A).The performance of the NG-Test VanB was validated by using a collection of 104 well-characterized enterococcal isolates grown on ChromID^®^ VRE, a media classically used for VRE screening from stool samples (9) or on MH for non-VRE isolates. All these isolates were characterized and provided by the French National Reference Center for VREs (12). This panel included 84 VREs (33 VanB-, 24 VanA-, 8 VanC1-, 12 VanC2-, 3 VanD-, 1VanE-, 1 VanG-, 1 VanL-, and 1 VanN-producers), 1 VanM-producing *E. coli* isolate, and 19 non-VRE isolates. Sixty-eight isolates were *E. faecium*, 20 *E. faecalis*, 8 *E. gallinarum* and 12 *E. casseliflavus* and one *E. coli* expressing VanM (12). All 33 VanB-VRE were detected in less than 15 min while no-cross reaction was observed with other acquired determinants (*i*.*e*., VanA, C1, C2, D, E, G, L, M, N) and non-VRE isolates. These results showed that the NG-Test VanB had sensitivity and specificity both equal to 100%. The limit of detection (LOD) was determined by using two VanB-*E. faecium* isolates grown on ChromID^®^VRE agar plates and corresponding serially-diluted bacterial suspensions. One hundred microliter of each dilution were mixed with 100 µL of EB-80. Serial dilutions were also plated on MH plates to determine the exact CFU/mL. The LOD was estimated at 0.95 × 10^7^ CFU per test. This LOD is two-log higher than that of the NG-Test VanA (4.9 10^5^ CFU/test) (12).

**Table 1.**
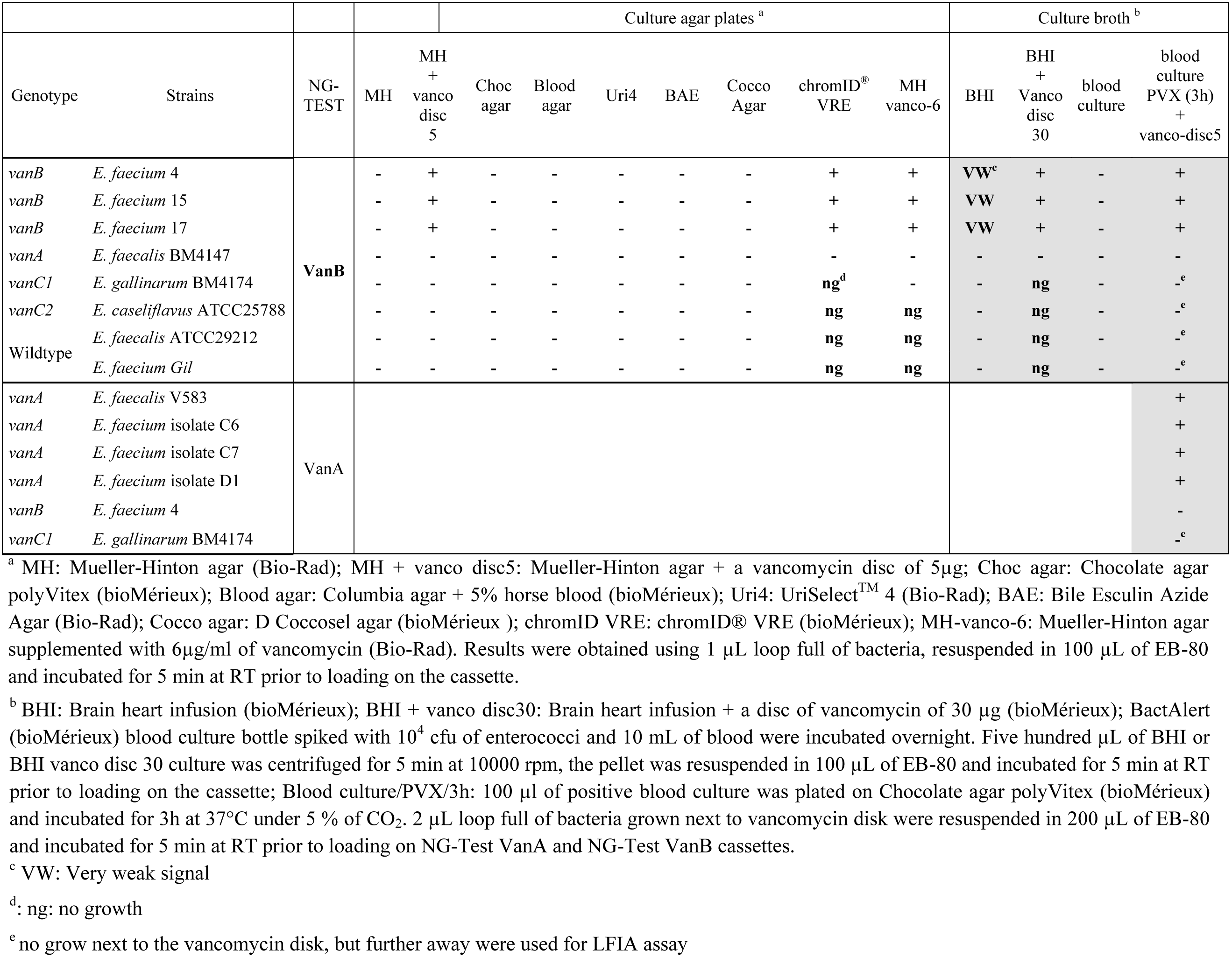
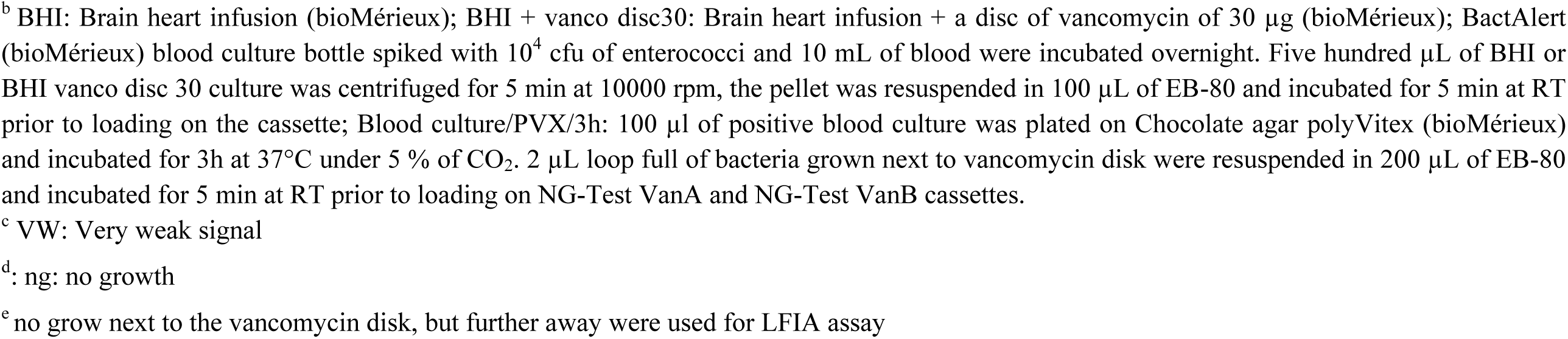
Results of the NG-Test VanB on colonies grown on different solid and liquid culture media and of the NG-Test VanA for isolates from blood culture.

**Figure 1.**
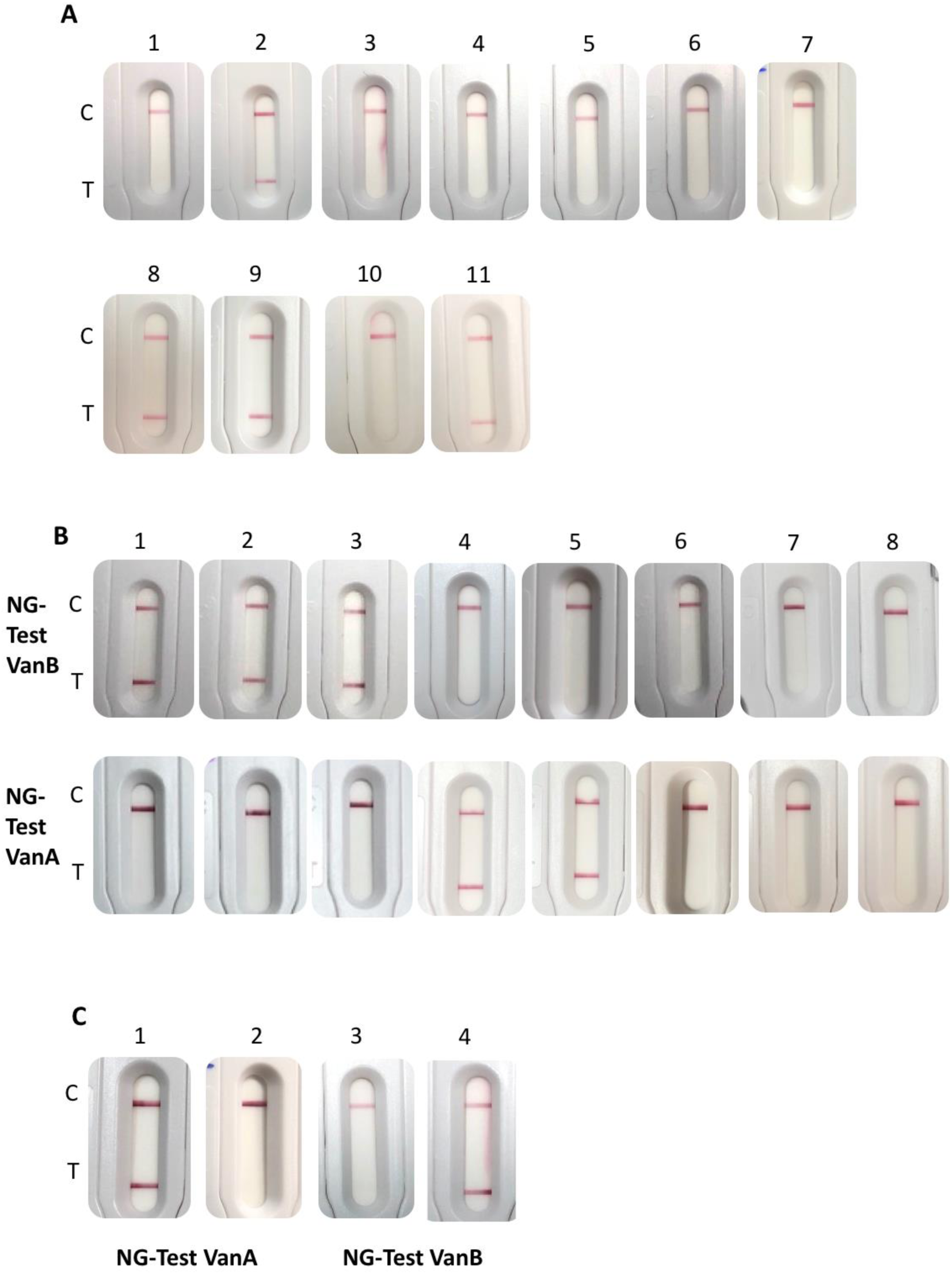
NG-Test VanB and VanA results. A. obtained with 1 µL loop full of *E. faecium* VanB grown on different agar plates: (1) Müller-Hinton (MH); (2) MH with a 5µg vancomycin disk; (3) Chocolate agar PolyVitex; (4) Columbia Agar + 5% of horse blood; (5) UriSelect4; (6) Bile esculin agar; (7) D-Coccosel agar; (8) ChromID^®^VRE; (9) MH supplemented with 6 mg/L of vancomycin; and with 500 µl of overnight grown *E. faecium* VanB (10) in Brain heart infusion (BHI); (11) and in BHI with a 30 µg disk of vancomycin. Spun down bacterial pellets were resuspended in 100 µL of EB-80 and incubated for 5 min at RT prior to loading on the cassette. B. obtained with 1 µL loop full of bacteria grown on ChromID^®^VRE, resuspended in 100 µL of EB-80 and incubated for 5 min at RT prior to loading on NG-Test VanB and NG-Test VanA cassettes. The tested bacteria were (1) *E. faecalis* VAN B; (2) *E. faecium* VAN B; (3) *E. faecium* VAN B; (4) *E. faecium* VanA isolate 12 (MIC Vancomycin/Teicoplanin 256/48 mg/L) (10); (5) *E. faecium* VanA isolate 2 (MIC Vancomycin/Teicoplanin 16/6 mg/L) (10); (6) *E. faecalis* ATCC29212; (7) *E. gallinarum* VanC1 BM4174; (8) *E. casseliflavus* VanC2 ATCC25788. **C**. obtained with spiked blood cultures. Blood cultures were spiked with 10^4^cfu of *E. faecium* VanA (1,3) and *E. faecium* VanB (2,4) and 10-ml of blood were incubated overnight. Subsequently, 100 µl of positive blood culture were plated on Chocolate agar polyVitex (bioMérieux) and incubated for 3h at 37°C under 5 % of CO_2_. 2 µL loop full of bacteria grown next to vancomycin disk were resuspended in 200 µL of EB-80 and incubated for 5 min at RT prior to loading on the NG-test VanA (1,2) and NG-Test VanB (3,4) cassettes.

**Figure 2:**
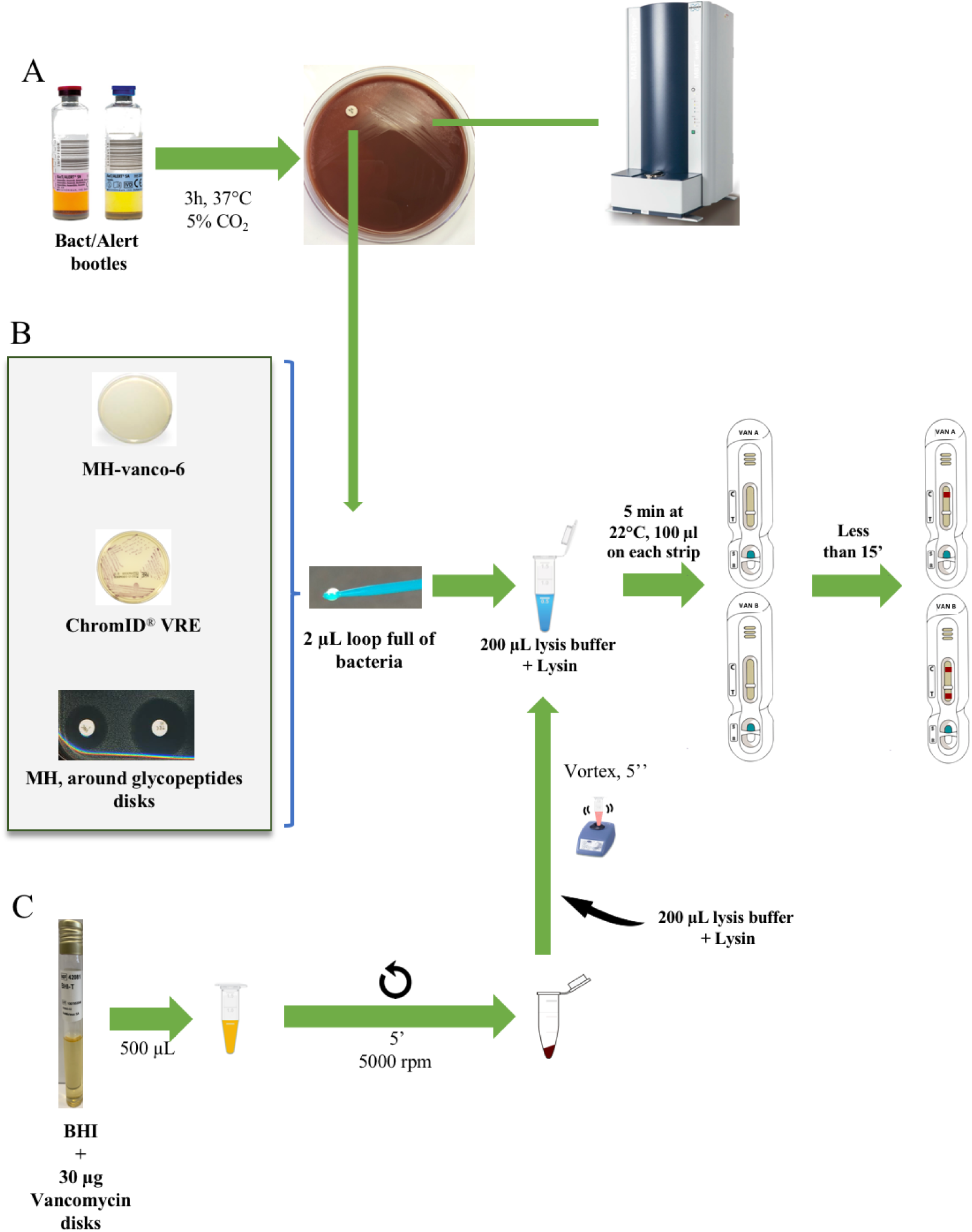
Experimental procedure. A. NG-Test VanA and NG-Test VanB on spiked blood cultures. Blood cultures were spiked with 104cfu of Enterococci and 10-ml of blood were incubated overnight. Subsequently, 100 µl of positive blood culture were plated on Chocolate agar polyVitex (bioMérieux) and incubated for 3h at 37°C under 5 % of CO2. 2 µL loop full of bacteria grown next to vancomycin disk were resuspended in 200 µL of EB-80 and incubated for 5 min at RT prior to loading 100 µL on the NG-test VanA and 100 µL on NG-Test VanB cassettes. Bacterial lawn was also used for MALDI TOF identification. B. from o/n grown bacteria on agar plates. MH-Vanco-6: Müller-Hinton supplemented with 6 µg/mL of vancomycin; ChromID^®^VRE (bioMérieux); and MH next to a 5µg vancomycin disc. 2 µL loop full of bacteria were resuspended in 200 µL of EB-80 and incubated for 5 min at RT prior to loading 100 µL on the NG-test VanA and NG-Test VanB cassettes C. From liquid broth. BHI: brain heart infusion and VRE broth were from bioMérieux. stands for centrifugation

Detection directly from positive blood cultures was also tested using 500 µl processed using the broth protocol (12) but it did not reveal a positive signal, as VanB production was not induced. We therefore implemented an optimized protocol. Positive blood cultures with spiked enterococci (3 VanB-VRE, 4 VanA-VRE, and 3 non-VanA and -B-VRE) were subcultured for three hours on Chocolate agar PolyViteX with a vancomycin disk (5 µg). Subsequently, colonies grown around the vancomycin disk were used for NG-TEST VanA and NG-Test VanB, while other colonies were used for MALDI-TOF mass spectrometry identification, as routinely performed in our laboratory. Using this protocol, enterococci were identified with MS scores > 1.8-2, thus allowing reliable identification at the species level and the presence of VanA or VanB could be evidenced using the two NG-Test strips, in less than 3h15 after blood culture was withdrawn from the automated incubator (BactAlert, bioMérieux) (Table 1). As our study was performed using small numbers of spiked blood cultures and not from real clinical samples, further prospective studies will be needed to estimate the performance of NG-Test VanA and VanB on blood cultures using our protocol.

In conclusion, the NG-Test VanB was able to detect VanB-VREs only when the bacteria were grown in the presence of 4-6 µg/mL of vancomycin. This is not a problem, as these tests may be used as confirmatory tests for detection of VanB, either from colonies growing on selective media containing vancomycin (such as ChromID VRE) or from a routine antibiogram next to vancomycin-containing disk. Direct detection from blood cultures is not possible, but as subculturing for three hours on Chocolate agar PolyViteX is routinely performed for MALDI-TOF mass spectrometry identification, addition of a 5-µg vancomycin disk could allow sufficient VanB induction for efficient and its reliable detection. The NG-Test VanB is easy to use, rapid, and does not require any specific equipment or skills while results are easy to read after 15 min of migration. As such, it can be easily implemented in the routine workflow of most clinical laboratories as confirmatory test of VanB-VRE. Together with the NG-Test VanA (12), they could complete/replace the already existing panel of tests available for confirmation of acquired vancomycin resistance in enterococci, especially from selective media (rectal screenings) or from antibiograms (infections), with a sensitivity and specificity both of 100%. The rapid detection in less than 15 minutes will result in more efficient management of carriers and infected patients. In the near future, the addition of emerging Van alleles, such as the VanD or VanM determinant, would allow detection of most resistance mechanisms encountered in VRE (4,16).

## Acknowledgments

We acknowledge NG Biotech for providing the NG-Test Van B and NG-Test VanA free of charge.

## Funding

This work was supported by the Assistance Publique-Hôpitaux de Paris (AP-HP), the University Paris-Saclay, the Laboratory of Excellence in Research on Medication and Innovative Therapeutics (LERMIT) supported by a grant from the French National Research Agency (ANR-10-LABX-33) and by the EIT health for the AMR-Detectool project.

## Transparency declarations

LB is a NG Biotech employee only involved in assay development but not in the validation process.

All other authors: None to declare.

## References

1. Uttley AH, Collins CH, Naidoo J, George RC. 1988. Vancomycin-resistant enterococci. Lancet 1:57–58.

2. Leclercq R, Derlot E, Duval J, Courvalin P. 1988. Plasmid-mediated resistance to vancomycin and teicoplanin in Enterococcus faecium. N Engl J Med 319:157–161.

3. WHO publishes list of bacteria for which new antibiotics are urgently needed. Available at: http://www.who.int/news-room/detail/27-02-2017-who-publishes-list-of-bacteria-for-which-new-antibiotics-are-urgently-needed. Accessed April 1, 2020.

4. Cattoir V, Leclercq R. 2013. Twenty-five years of shared life with vancomycin-resistant enterococci: is it time to divorce? J Antimicrob Chemother 68:731–742.

5. Shadel BN, Puzniak LA, Gillespie KN, Lawrence SJ, Kollef M, Mundy LM. 2006. Surveillance for Vancomycin-Resistant Enterococci: Type, Rates, Costs, and Implications. Infect Control & Hosp Epidemiol 27:1068–1075.

6. Bender JK, Cattoir V, Hegstad K, Sadowy E, Coque TM, Westh H, Hammerum AM, Schaffer K, Burns K, Murchan S, Novais C, Freitas AR, Peixe L, Del Grosso M, Pantosti A, Werner G. 2018. Update on prevalence and mechanisms of resistance to linezolid, tigecycline and daptomycin in enterococci in Europe: Towards a common nomenclature. Drug Resist Updates 40:25–39.

7. Arthur M, Depardieu F, Reynolds P, Courvalin P. 1996. Quantitative analysis of the metabolism of soluble cytoplasmic peptidoglycan precursors of glycopeptide-resistant enterococci. Mol Microbiol 21:33–44.

8. Papadimitriou-Olivgeris M, Filippidou S, Kolonitsiou F, Drougka E, Koutsileou K, Fligou F, Dodou V, Sarrou S, Marangos M, Vantarakis A, Anastassiou ED, Petinaki E, Spiliopoulou I. 2016. Pitfalls in the identification of Enterococcus species and the detection of vanA and vanB genes. Lett Appl Microbiol 63:189–195.

9. Cuzon G, Naas T, Fortineau N, Nordmann P. 2008. Novel chromogenic medium for detection of vancomycin-resistant Enterococcus faecium and Enterococcus faecalis. J Clin Microbiol 46: 2442–4.

10. Naas T, Fortineau N, Snanoudj R, Spicq C, Durrbach A, Nordmann P. 2005. First nosocomial outbreak of vancomycin-resistant Enterococcus faecium expressing a VanD-like phenotype associated with a vanA genotype. J Clin Microbiol 43: 3642–9.

11. Rasoanandrasana S, Decousser J-W, Cattoir V, Berçot B, Domrane C, Fihman V, Grillon A, Lesenne A, Raskine L, Cambau E, Jacquier H. 2017. Use of ESwab in the Xpert® vanA/vanB PCR assay. Eur J Clin Microbiol Infect Dis 2017 36: 755–6.

12. Oueslati S, Volland H, Cattoir V, Bernabeu S, Girlich D, Dulac D, Plaisance M, Laroche M, Dortet L, Simon S, Naas T. 2020. Development and validation of a lateral flow immunoassay for rapid detection of VanA-producing enterococci, J Antimicrob Chemother, dkaa413, https://doi.org/10.1093/jac/dkaa413

13. Boutal H, Naas T, Devilliers K, Creton E, Cotellon G, Plaisance M, Oueslati S, Dortet L, Jousset A, Simon S, Naas T, Volland H. 2017. Development and Validation of a Lateral Flow Immunoassay for Rapid Detection of NDM-Producing Enterobacteriaceae. J Clin Microbiol 55: 2018–29.

14. Bernabeu S, Ratnam KC, Boutal H, Gonzalez C, Vogel A, Devilliers K, Plaisance M, Oueslati S, Malhotra-Kumar S, Dortet L, Fortineau N, Simon S, Volland H, Naas T. 2020. A Lateral Flow Immunoassay for the Rapid Identification of CTX-M-Producing Enterobacterales from Culture Plates and Positive Blood Cultures. Diagnostics (Basel). 10 (10):764. doi: 10.3390/diagnostics10100764.

15. Yoong P, Schuch R, Nelson D, Fischetti VA. 2004. Identification of a broadly active phage lytic enzyme with lethal activity against antibiotic-resistant Enterococcus faecalis and Enterococcus faecium. J Bacteriol; 186: 4808–12.

16. Sun L, Qu T, Wang D, et al. Characterization of vanM carrying clinical Enterococcus isolates and diversity of the suppressed vanM gene cluster. Infect Genet Evol 2019; 68: 145–52.

